# Understanding Cu^+2^ binding with DNA: A molecular dynamics study comparing Cu^2+^ and Mg^2+^ binding to the Dickerson DNA

**DOI:** 10.1101/2024.07.04.602035

**Authors:** Angad Sharma, Hari O. S. Yadav, Pradipta Bandyopadhyay

## Abstract

Cu^2+^ ions led DNA damage by reactive oxygen species (ROS) is widely known biological phenomena. The ionic radii of Cu^2+^ and Mg^2+^ being similar, the binding of Cu^2+^ ions to DNA is expected to be similar to that of the Mg^2+^ ions. However, little is known how Cu^2+^ ions bind in different parts (phosphate, major and minor grooves) of a double-strand (ds) DNA, especially at atomic level. In the present study, we employ molecular dynamic (MD) simulations to investigate the binding of Cu^2+^ ions with the Dickerson DNA, a B-type dodecamer double stranded (ds) DNA. The binding characteristics of Cu^2+^ and Mg^2+^ ions with this dsDNA are compared to get an insight into the differences and similarities in binding behavior of both ions. Unlike Mg^2+^ ions, the first hydration shell of Cu^2+^ is found to be labile, thus it shows both direct and indirect binding with the dsDNA, i.e., binding through displacement of water from the hydration shell or through the hydration shell. Though the binding propensity of Cu^2+^ ions with dsDNA is observed relatively stronger, the binding order to phosphates, major groove, and minor groove is found qualitatively similar (phosphates > major groove > minor groove) for both ions. The study gives a deep understanding of Cu^2+^ binding to DNA, which could be helpful in rationalizing the Cu^2+^ led ROS-mediated DNA damage.

## INTRODUCTION

Deoxyribonucleic acid (DNA) and ribonucleic acid (RNA), two forms of nucleic acids, are extremely significant macromolecules in biology that carry the genetic code of all living things and provide necessary instructions for protein synthesis^1–3^. Due to the negative charge on the oxygen of the phosphate groups, these nucleic acids exist as the polyanionic macromolecules.^4^ Thereby, they are always surrounded by a thick layer of counterions, which is often termed as the “ion atmosphere”. This ionic atmosphere is mostly composed of intracellular ions (Na^+^, K^+^, Mg^2+^, Ca^2+^, etc.)^5^, but other metal ions can also interact with the DNA and RNA depending upon the toxicity level of the metal ions in the body^6^. These counterions balance out the negative charges of nucleic acids and thus provide them with structural stability and function^7,8^. Additionally, they actively participate in modulating genome packaging^9^, DNA and RNA folding^10^, enzyme activity^11^, and serve as the indispensable mediators in the DNA-protein interaction^12^.

Copper is an essential trace element for human body^13^. Its antimicrobial property is known since Egyptian era^14,15^. It was used to cure wounds and clean water^16^. When antibiotics were not available, many diseases such as tuberculosis, syphilis, skin infections used to treat with copper^17^. Its deficiency in the body can affect several biological functions such as tissue synthesis, cellular respiration, gene expression, iron metabolism, free radical defense as well as the normal operation of heart and immunes^18–20^. Regardless of many therapeutic properties, excess copper can pose a serious health risk. The excess can cause several hereditary syndromes such as Wilson disease, idiopathic copper toxicosis etc^21^. Copper ions (Cu^2+^) can strongly bind with the double-strand DNA (dsDNA) and affect its double helical structure. As a result, the inherent biological function of dsDNA can be modulated^22^. In fact, the studies show that Cu^2+^ can cause permanent damage in the dsDNA by producing the reactive oxygen species^23^. Cu^2+^ ions bound to dsDNA can react with H_2_O_2_ and ascorbic acid present in the body and generate hydroxyl free radicals, which can immediately attack dsDNA bases in a specific manner and result in damage. The damage includes breaking of DNA strands, disruption of the hydrogen-bond network of bases as well as other changes in the DNA bases. These changes can lead to mutations and potentially contribute to diseases like cancer and hereditary syndromes.^24,25^

Several experimental studies^26–36^ have been performed with the aim of identifying specific binding sites in the dsDNA for Cu^2+^ ions. It was suggested that the binding of Cu^2+^ ion to DNA is highly cooperative in nature, i.e. the binding happens with DNA denaturation or at least with local deformation of the double helix^30,35^. It was found that Cu^2+^ has the highest affinity among the divalent metal cations that bind to the DNA^35^. At very low concentration, Cu^2+^ ions bind non-specifically with the phosphate groups, but as the concentration increases, they begin to bind with the DNA bases (e.g. with O and N atoms of DNA bases) too, in fact with much higher affinity than for the phosphate groups^34^. It was shown that Cu^2+^ ions mainly bind with guanine (G) and cytosine (C) bases^28^. In guanine base, Cu^2+^ can coordinate with N7 and O6 atoms and in cytosine it can coordinate with N3 and O2 atoms (see figure 1 for the definitions for these atoms). Considering these facts, various binding models for Cu^2+^ ions to DNA bases were hypothesized^28,29^. Richard et al^28^ proposed two binding models for Cu^2+^ ions to DNA bases: (1) a “sandwich” structure of Cu^2+^ ion between to G-C pairs, and (2) a chelate between a phosphate group and a nitrogen atom of bases. The nitrogen atom could be of N7 of guanine or N3 of cytosine as discussed above. Cu^2+^ can coordinate with N3 of adenine too but it can happen only at higher temperature, where the thermal energy higher than the room temperature can cause opening of adenine-thymine (A-T) base pair^28^. Out of these two models, the latter is the most widely accepted model for Cu^2+^ ion binding to DNA.

**Figure 1.**
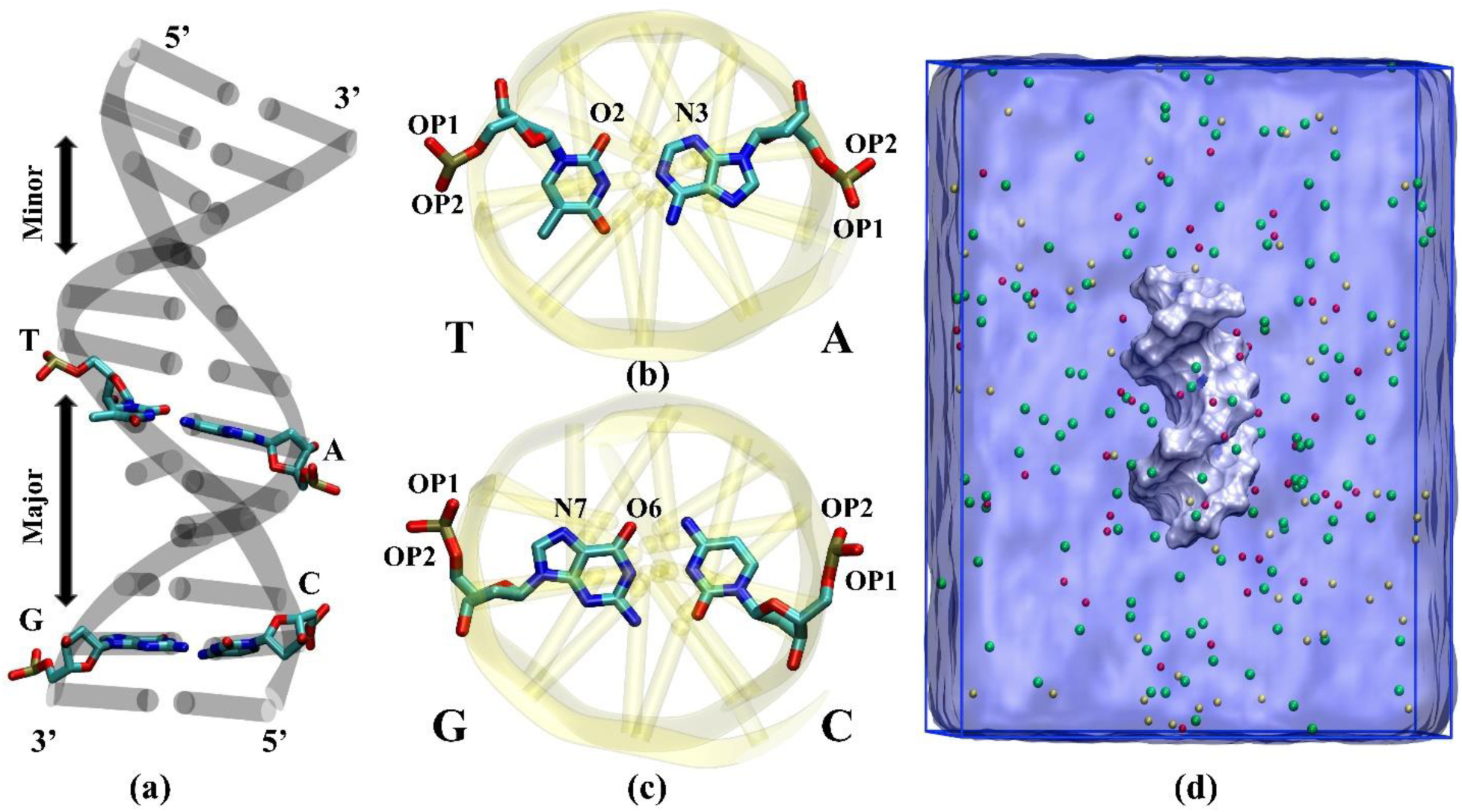
(a) A double stranded DNA structure showing major and minor grooves, (b) and (c) shows the atoms used in the analysis (d) Hydrated structure of the DNA. The silver surface represents the DNA; the continuum light-blue phase is water; the red, yellow, and green beads are Cu^2+^, Na^+^, and Cl^-^ ions, respectively.

Interaction of DNA and ions was also studied with several theoretical approaches such as Manning condensation model^37^, Poisson-Boltzmann (PB) theory^38^, three-dimensional reference interaction site models (3D-RISM)^39^ but these methods do not fully describe the DNA-ion binding, especially the binding of multi-valent cations^40,41^. The reason for this is the many simplifying assumptions made by these theories. For example, Manning model neglects the atomic-level properties of ions and solvent as well as the charge-density variations along the DNA backbone^4^. PB theory does not account the ion size or ion-ion correlation and treat their interactions in an averaged fashion. Furthermore, it considers the water molecules and the interior of the DNA as media with uniform dielectric constants. The results coming from 3D-RISM may depend significantly on the specific closure relationship used in the calculation. A few quantum-mechanical (QM) studies are also available on the binding of Cu^2+^ ions to DNA^42–44^. These studies show that the highest affinity of Cu^2+^ ion among DNA bases is to guanine and the favorable binding sites are N7 and O6 atoms.

However, all these previous studies mostly focused on predicting the favorable binding sites for Cu^2+^ ions in DNA, structural changes of DNA, and properties of DNA-Cu^2+^ adducts. It is still not well understood in what proportions Cu^2+^ ions bind to different parts of the DNA, i.e., on the phosphate, major, and minor grooves. In other words, the specific localization of Cu^2+^ ions on the phosphate group, within the major and minor grooves is not understood yet, especially at the atomic resolution. In general, an ion can bind with the DNA in the partially dehydrated or fully hydrated form, where the former is termed as a “direct” and the latter as an “indirect” interaction^6^. Molecular details of the hydration state of Cu^2+^ ion bound to DNA or changes in the hydration state as it approaches to the DNA from bulk solvent region are still obscure and need an in-depth investigation. As a result of such obscurities, the binding action mechanism of Cu^2+^ ion to DNA is still not clear. Extensive MD simulation studies of DNA interactions with alkali and alkaline earth metal ions exist^45–49^, but interaction studies of DNA with transition metal ions, especially Cu2+, are not well available.

In the present study, the binding of Cu^2+^ and Mg^2+^ ions on the phosphate group, major, and minor grooves of a Dickerson–Drew (DD) dodecamer sequence (d-[CGCGAATTCGCG]_2_) is investigated using the molecular dynamics (MD) simulations. DD sequence is a prototypic B-type double-strand DNA, whose structure as well as dynamics have been extensively studied with both experiment^50–52^ and simulation^53–55^. The interaction between Mg^2+^ ion and DNA is included in the study as a reference system because its interaction with DNA is relatively better understood than that of other divalent cations. DNA-ion binding is analyzed through structural metrices and free energy calculation. The differences and similarities in the binding behavior of Cu^2+^ and Mg^2+^ are thoroughly researched and rationalized. To the best of our knowledge, this is the first MD simulation-based study of the binding of Cu^2+^ ion to DNA. It is believed that this study would enhance our understanding of DNA and Cu^2+^ interaction and aid in experiments.

## MATERIALS AND METHODS

### Molecular Dynamics Simulation

All MD simulations were performed using the AMBER16 MD package^56,57^ with GPU acceleration for non-bonded interactions and applying the periodic boundary conditions (PBC) in all directions. The initial structure of the Dickerson–Drew dodecamer sequence (PDB ID: 1bna) was taken for protein data bank^50^. In the native state, this DNA sequence had a major and minor groove, and 22 negative charges attached to phosphate groups. For solvated initial configuration, DNA structure was put at the center of the octahedral simulation box and then box randomly filled with water molecules without overlaps of atoms. The simulation box (dimension was 88.6*92.8*109.8 Angstrom^3^) was taken sufficiently large to avoid the spurious interactions of DNA with its periodic images. Negative charges of DNA were neutralized with Na^+^ ions and then an additional 0.10 M NaCl salt was added in the solution to mimic the physiological conditions. After this, 0.15 M CuCl_2_ salt was introduced in the simulation cell to study the interaction of Cu^2+^ions with DNA. A depiction of the simulation box is shown in Fig 1. The reference system, i.e., the interaction of DNA with Mg^2+^ ions, was studied at the same concentration and its initial hydrated configuration was prepared in the same manner. All initial configurations were generated using LEaP tool of AMBER and visualized using VMD package^58^.

DNA was modeled with parmBSC1 force field^59^. The interactions of monovalent ions were defined using Jong-Cheetham parameters^60^ and interactions of divalent Cu^2+^ and Mg^2+^ ions were denoted by Li-Merz parameters^61^. Water molecules were described with rigid TIP3P model^62^. Note that the employed interaction parameters of ions are already present in the BSC1 library and the cross interaction of dissimilar pairs are determined using the Lorentz-Berthelot rules^63^. Rigid bonds and rigid angles of water molecules or elsewhere were dealt with SHAKE algorithm^64^. The short-range Lennard-Jones interaction was truncated at 10 Å and long-range electrostatic interactions were calculated using the particle mesh Ewald algorithm^65^. The production MD were carried out in isothermal-isobaric (NPT) ensemble. Simulation temperature was controlled using the Langevin thermostat^66^ with collision frequency of 1 ps^-1^ and pressure was fixed with Berendsen barostat^67^ using a time constant of 2 ps. Newtonian equation of atoms were integrated using the velocity-Verlet algorithm^68^ with a timestep of 2 fs.

Minimization of each system was carried out in two phases. In the first step, minimization was performed by imposing a harmonic restraint on the positions of each of the heavy atoms of the DNA with a harmonic force constant of 500 kcal mol^-1^. Å^-2^ for 1500 steps. Minimization of the first 1000 steps was done using the steepest descent method^58^ and the next 500 steps were done using the conjugate gradient algorithm^63^. In the second phase, minimization of the entire system was performed for 2500 steps using the conjugate gradient method without restraints. After energy minimization, solvated DNA-system was gradually heated from 0 K to 300 K in the NVT ensemble with a heating rate of 6 K ps^-1^. After the temperature reached to 300 K, an equilibration MD in the NPT ensemble was performed for 1 ns and then a production MD for 1 microsecond with frequency of the trajectory at every 20 ps. Three independent simulations were performed for each system with different initial configurations for better average and error estimation.

Binding free energy, i.e. the potential of mean force (PMF) between an electronegative site in the DNA and an ion (Cu^2+^ or Mg^2+^) as a function of distance was calculated using the umbrella sampling method^69^. Two types of force constants of harmonic umbrella potential (bias) were used in the study. For binding of ions to phosphate group (i.e. to OP1 or OP2), the force constant was chosen to be 150 kcal mol Å^-2^. For binding sites in bases, it was taken 100 kcal mol Å^-2^. The ion from the binding site of interest was initially placed at 6 Å. Umbrella sampling was carried out along the reaction coordinate (i.e. the distance between the ion and binding site) with a window spacing of 0.1 Å. For each window, an equilibration was performed for 1 ns, followed by a production MD for 10 ns. The statistical data (i.e., the force and distance) in each window were dumped at every 50 steps. PMF was extracted from these data using the weighted histogram analysis method (WHAM)^70^.

## RESULTS AND DISCUSSION

### Hydration Structure of Cu^2+^ and Mg^2+^ Ions

Hydration shell of an ion can play a vital role in the DNA-ion interaction. For direct binding to a site in DNA, an ion must undergo partial dehydration. To comprehend such scenarios, we first investigated the hydration shell structure of Cu^2+^ and Mg^2+^ ions in bulk water. Figure 2(a) shows the radial distribution function, *g*(*r*), and coordination number, *N_c_*(*r*), of oxygen atoms of water around Cu^2+^ and Mg^2+^ ions. The *g*(*r*) provides a quantitative measure of the packing structure and probability of finding water molecules around ions, while *N_c_*(*r*) bestows the variation of the number of water molecules as a function of distance, *r*, from ions. For both ions, two distinct peaks appear in the *g*(*r*), which correspond to the first and second hydration shells of the ions. The result shows that the first peak of *g*(*r*) for Mg^2+^ ion is narrower and its height is higher than that for Cu^2+^ ion. The observation suggests that the first hydration shell of the Mg^2+^ ion is more well-structured than that of the Cu^2+^ ion. The height of the second peak of *g*(*r*) is almost the same for both ions but is broader than the first peak indicating the diffused nature of the second hydration shell. For both ions, the *N_c_*(*r*) of the first hydration shell is six, which infers that the first hydration shell of both ions are formed of six water molecules. This is also evident in the visual snapshots of the first hydration shell of the ions shown in Figures 2(b) and 2(c). The *N_c_*(*r*) profiles demonstrate that the coordination number of Cu^2+^ ion in the second hydration shell is slightly higher than that of Mg^2+^ ions and constitutes of 12 – 14 water molecules.

**Figure 2.**
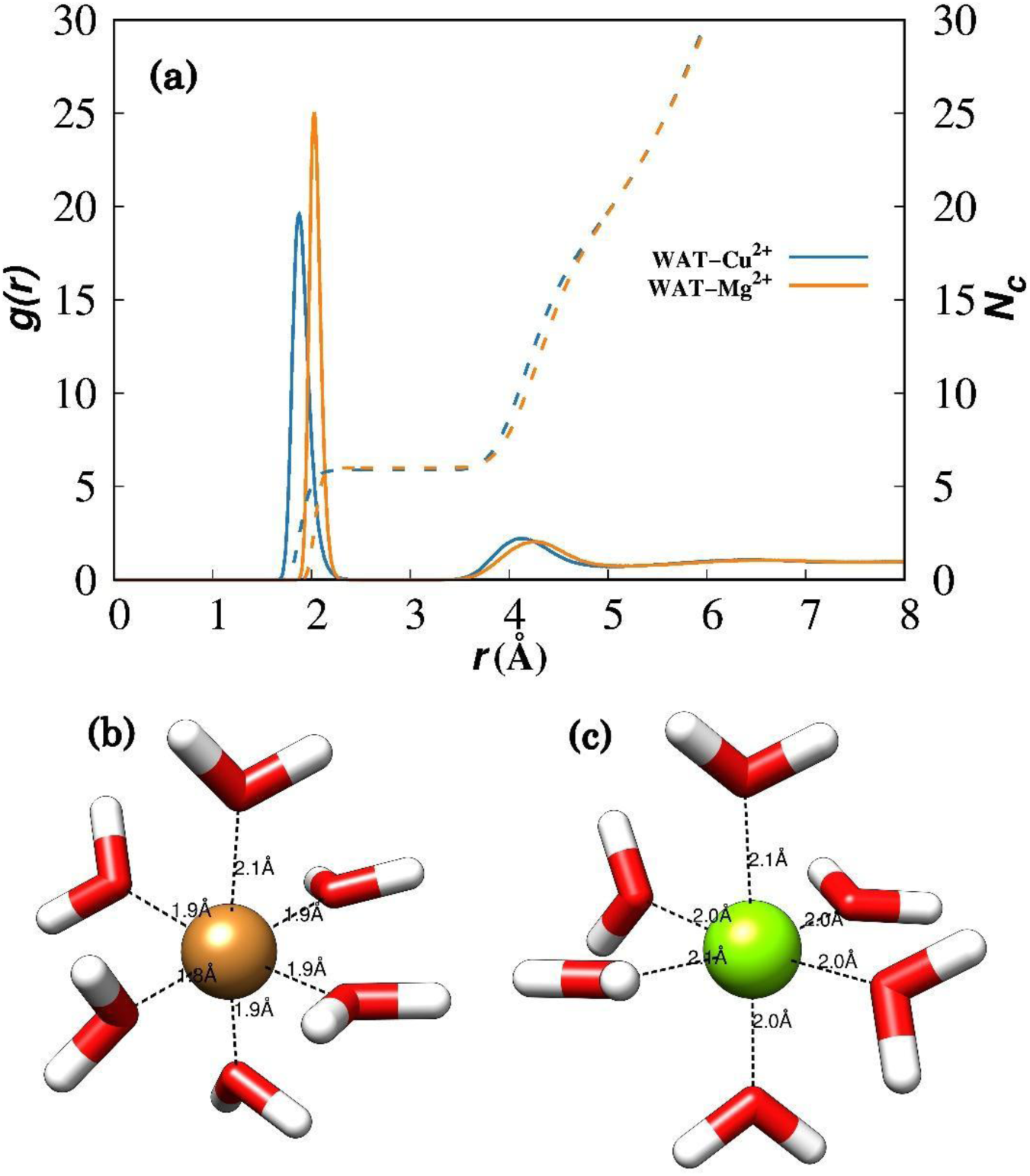
(a) Radial distribution function (solid line), *g*(*r*), and coordination number (dotted line), *N_c_*, of oxygen atoms of water around Cu^2+^ and Mg^2+^ ions in bulk water, where *r* is the distance between the oxygen atom of water and ion. The data of *N_c_* is shown on y2-axis. Subfigure (b) and (c) shows the visual of the first hydration shell of Cu^2+^ (brown bead) and Mg^2+^ (green bead) ions, respectively. Water molecules are displayed in licorice model, with the red portion representing the oxygen atom and the white portions representing the hydrogen atoms.

We also examined the coordination number of ions present around the DNA as the coordination shells of these ions may differ in the presence of DNA and monovalent ions (Na^+^ and Cl^-^) used in the simulation. Figure 3 shows the probability distribution of *c*oordination number, *P*(*N_c_*), of Cu^2+^ and Mg^2+^ ions, which are within 5 Å of DNA. Here, *N_c_* correspond to the first hydration shell and calculated as the number of water oxygen or Cl^-^ lying within 2.5 Å of the cations. The cutoff distance (2.5 Å) was chosen from *g*(*r*) of ion and bulk water, which clearly demonstrate that the first hydration shell lies within this radius (see Figure 2). One can notice that the Mg^2+^ ion has only one peak in the *P*(*N_c_*) centered on 6. This suggests that the coordination number of Mg^2+^ ion near DNA is similar to its coordination number in bulk water. In other words, the coordination number of Mg2+ ion does not change near the DNA and it interacts with the DNA in fully hydrated form. On the contrary, *P*(*N_c_*) of Cu2+ ion is broad and spanned over 3 to 6. This suggests that unlike the Mg^2+^ ion, the hydration shell of the Cu2^+^ ion is flexible (despite its more favorable hydration free energy (see Fig. S1)) and it interacts with DNA through different coordination shells of water molecules. Note that the coordination shell of Cu^2+^ ion lying within 5 Å of the DNA has Cl^-^ ions also along with the water; however, the probability of observing Cl^-^ ions appears quite low. Its presence in the coordination shell (1 or 2 in the numbers) is observed only for 5-10% of Cu^2+^ ions.

**Figure 3.**
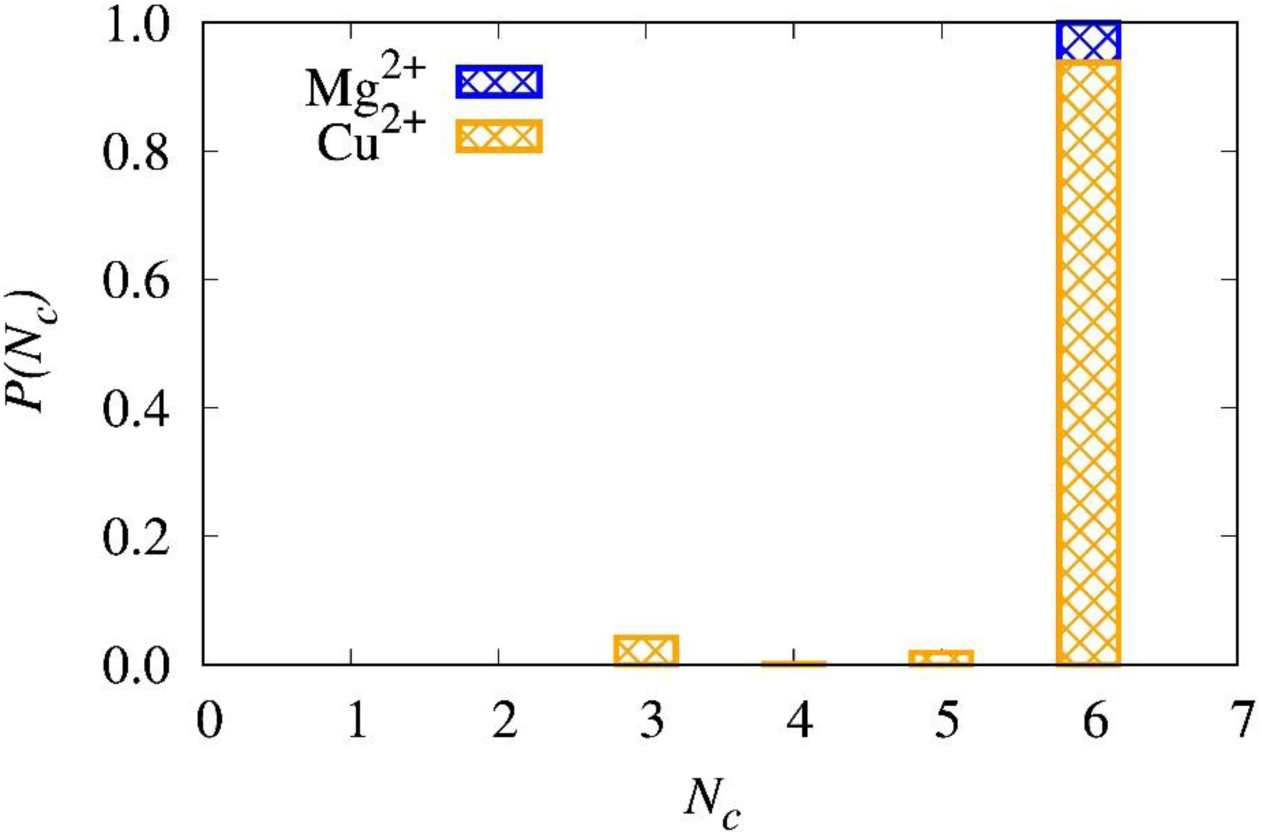
Probability distribution of coordination number, *P*(*N_c_*), of Cu2+ and Mg2+ ions lying within 5 Å of DNA.

### Binding Interaction of Metal Ions with DNA

#### Analysis of radial distribution functions

First, we consider ion binding to the phosphate groups. Each phosphate group possesses four oxygen atoms. Of these four, two are involved in the phosphodiester bonds. The other two lie in the outer layer exposed to the solvent. Metal ions can bind to both of these oxygens through electrostatic attraction. To get an insight into this interaction, we calculated the radial distribution function, *g*(*r*), of metal ions with OP1 and OP2 atoms separately and then averaged the results. The *g*(*r*) provides the probability of finding metal ions as a function of distance from phosphate oxygen atoms. It can be easily seen from Fig 2(a) that *g*(*r*) is highly structured (with multiple peaks) suggesting a layered structure of ions around DNA. In the *g*(*r*) between the Cu^2+^ and phosphate oxygens, the first prominent peak appears below 2.5 Å. This distance is less than the diameter of a water molecules, therefore the peak below 2.5 Å suggests that some Cu^2+^ ions are directly bonded with phosphate oxygen, i.e. they are interaction with phosphate oxygens in partially dehydrated form. It is even evident in the visual snapshot shown in Fig 4(b). The peaks visible at ≈ 4 and 6 Å corresponds to Cu^2+^ ions indirectly interacting with the phosphate oxygens, i.e. the Cu^2+^ ions interacting with OP1 and OP2 with complete hydration shell. The peak below 2.5 Å is missing in *g*(*r*) of Mg^2+^, which suggests that the Mg^2+^ ions have only indirect interaction with the phosphate groups as clear in the snapshot shown in Fig. 4(c). Further, the height of the peak appearing at ≈ 4 Å is lower than that of Cu^2+^ ions. This observation suggests that the interaction of Cu^2+^ ions with the phosphate group is more favorable than that of Mg^2+^ ions. In comparison to Mg^2+^ ions, Cu^2+^ ions show both direct and indirect interactions with the phosphate group because its hydration layer is labile as explained in the previous section.

**Figure 4.**
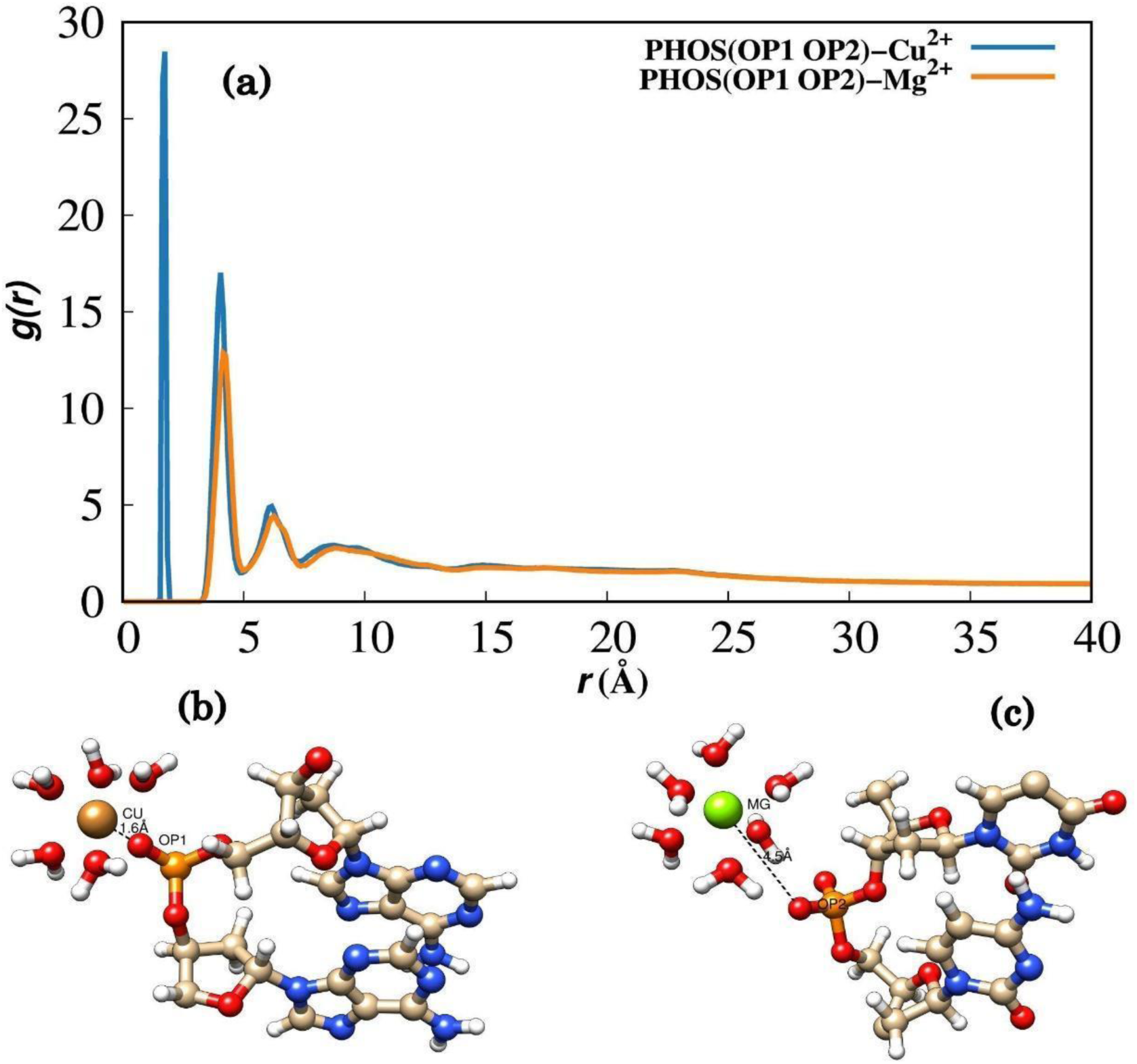
(a) The radial distribution function, *g*(*r*), of Cu^2+^ and Mg^2+^ with OP1 and OP2 sites of the phosphate groups of the DNA. The visuals of binding mode of (b) Cu^2+^ and, (c) Mg^2+^ ions with the oxygen of phosphate group.

DNA bases possess many electronegative (O and N) atoms, which can interact with metal ions. To understand the binding behavior of ions in DNA grooves, As depicted in Figure 5, we calculated the *g*(*r*) of metal ions with the N7 and O6 atoms in the major groove, and with the N3 and O2 atoms in the minor groove. The result clearly shows that the first peak in all cases appear at ≈ 4 Å, which suggests that there is no direct interaction of metal ions with DNA base atoms. To put it differently, there is no chelate formation of metal ions with DNA base atoms. This is contrary to the experiment for Cu^2+^ ions. Experimental studies suggest that the binding or chelate formation of Cu^2+^ is highly cooperative in nature i.e., it happens with DNA denaturation. However, in the present study, the RMSD data suggest that the structural change in DNA is negligible (see Fig. S2). Therefore, this could be a reason for indirect interaction of metal ions in the major and minor grooves. The height of first peak of *g*(*r*) of Mg^2+^ ions in the major groove for both N7 and O6 atoms is higher than that for Cu^2+^ ions. This implies that Mg^2+^ ions have more favorable interaction with DNA base atoms in the major groove than Cu^2+^ ions. This could be a dominant effect of the slightly lower number of water molecules in the hydration shell of Mg^2+^ ions, especially in the second hydration shell (see Fig 2). In general, the height of first peak of *g*(*r*) for O6 atoms is higher than that for N7 atoms. This is because the oxygen atom is more electronegative than the nitrogen atoms and has a smaller radius. Moreover, the dominant effect of direct electrostatic interactions, which are in general more favorable for Cu^2+^ ions (see Fig S3). In the minor groove, the binding of both the divalent ions with electronegative atoms i.e. N3 and O2 are negligible. This suggests these hydrated divalent ions are not able to enter in the minor groove. The interaction of these ions with N3 and O2 is happening indirectly with the hydration layer. The peak height of *g*(*r*) in the major groove is higher than in the minor groove, thus Fig. 4 and 5 in conjunction suggest that the binding affinity of ions is highest for phosphate groups, then in the major groove and lowest in the minor groove. The occupancy analysis shown in Fig. S4 provides similar findings. An ion is considered bound if its distance from the DNA is approximately 5 to 6 Å^49^. The impact of background Na^+^ and Cl^-^ ions was also analyzed by examining the number of ions surrounding DNA as a function of distance (*r*) in Figure S5. The analysis shows that Cu²⁺ ions are more abundant than Mg²⁺ and Na⁺ ions within 5 Å of the DNA. The Na⁺ ion count remains unchanged while there were more Cl⁻ ions present when DNA interacted with Cu²⁺ ions compared to when it interacted with Mg²⁺ ions.

**Figure 5.**
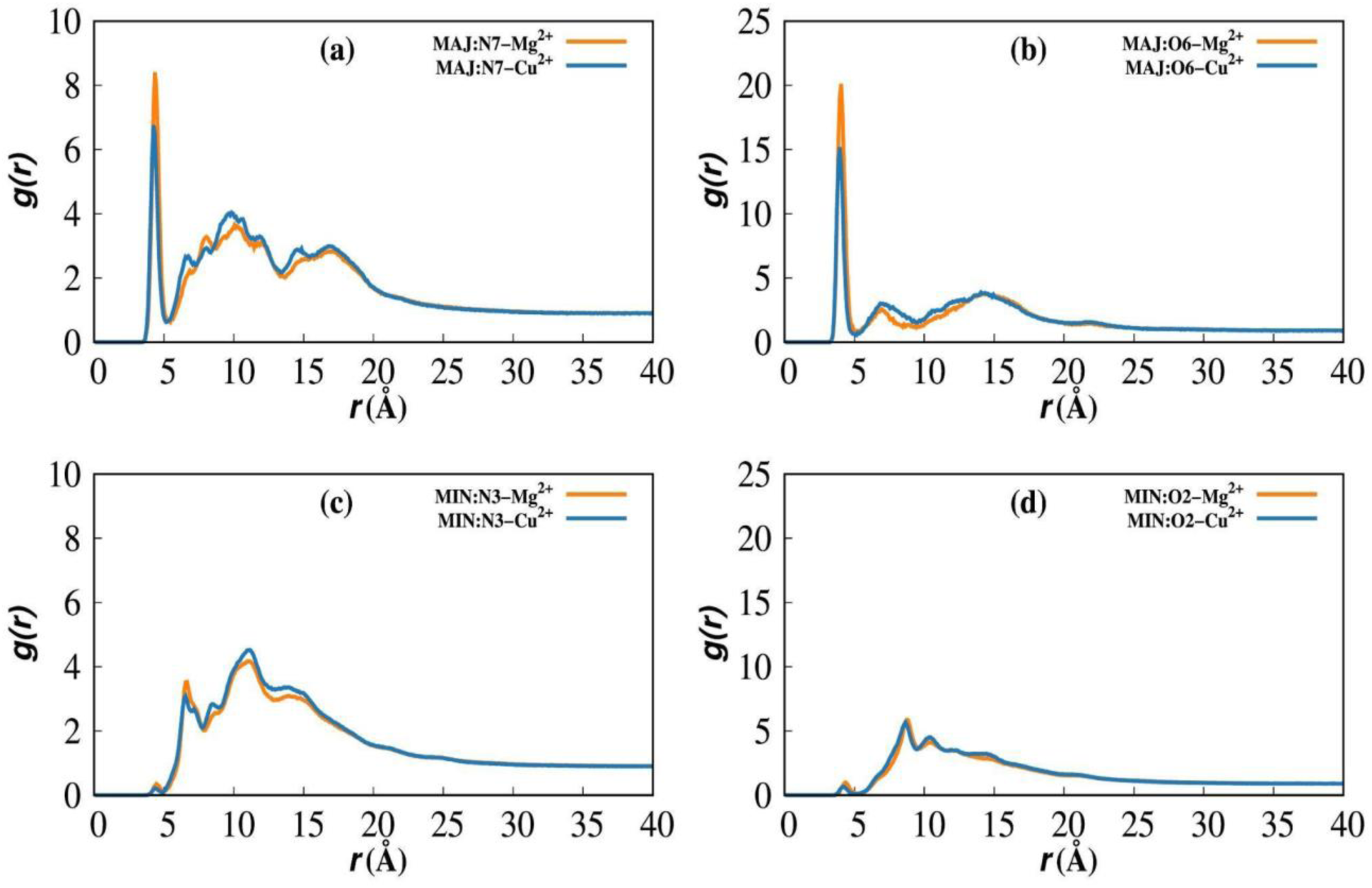
g(r) between the metal ions (Cu²⁺ and Mg²⁺) and electronegative atoms of DNA bases in the major and minor grooves. (a) and (b) show interactions with N7 and O6 atoms in the major groove, while (c) and (d) depict interactions with N3 and O2 atoms in the minor groove.

#### Potential of Mean Force (PMF) calculations

Ion binding was further investigated in energetic terms by calculating the potential of mean force, *F*(*r*), of the ion at a given atomic site in the DNA, *r* being the distance between the ion and the DNA site. Figure 6 shows the *F*(*r*) of a Cu^2+^ and Mg^2+^ ion with a OP2 atom of the phosphate group in the third nucleotide (cytosine) in the DNA sequence. This site was specifically chosen in the *F*(*r*) calculation because it shows direct binding with the Cu^2+^ ion in the unbiased MD simulations. From figure 6, it can be noticed that above 3 Å, one can notice that there is negligible difference in the free energy. In this region, both Cu^2+^ and Mg^2+^ ions are interacting with the DNA or given binding site through the complete first hydration shell, which is discussed in detail later. Below 3 Å, *F*(*r*) of both ions shows a maximum, i.e., positive energy barrier, then decreases rapidly and reaches a minimum. This decrease is due to partial dehydration of the first hydration shell, i.e. at least one water molecule is lost from the first hydration. Since the ion is less screened by water molecules, it interacts more favorably with the binding site through electrostatic interactions. The minimum for the Cu^2+^ ion is deeper than that for the Mg^2+^ ion because the direct electrostatic interaction of the Cu^2+^ ion with DNA is in general stronger than that of the Mg^2+^ ion. Going further below this minimum, the van der Waals repulsion between the ion and the binding site sets in and causes an increase in *F*(*r*). It is important to note that the positive free energy barrier (≈ +12.5 kcal/mol) appearing below 3 Å is more than twice the minimum (≈ -5 kcal/mol) seen around 2 Å in magnitude, while it is comparable (both ≈ 10 kcal/mol in magnitude) for the Cu^2+^ ion. This may be the key reason why only Cu^2+^ ions show direct binding with the DNA and not Mg^2+^ ions. Conclusively, the results show that the interaction of Cu^2+^ ion with the DNA phosphate group is more favorable than that of Mg^2+^ ion, which is consistent with the prediction of g(r).

**Figure 6.**
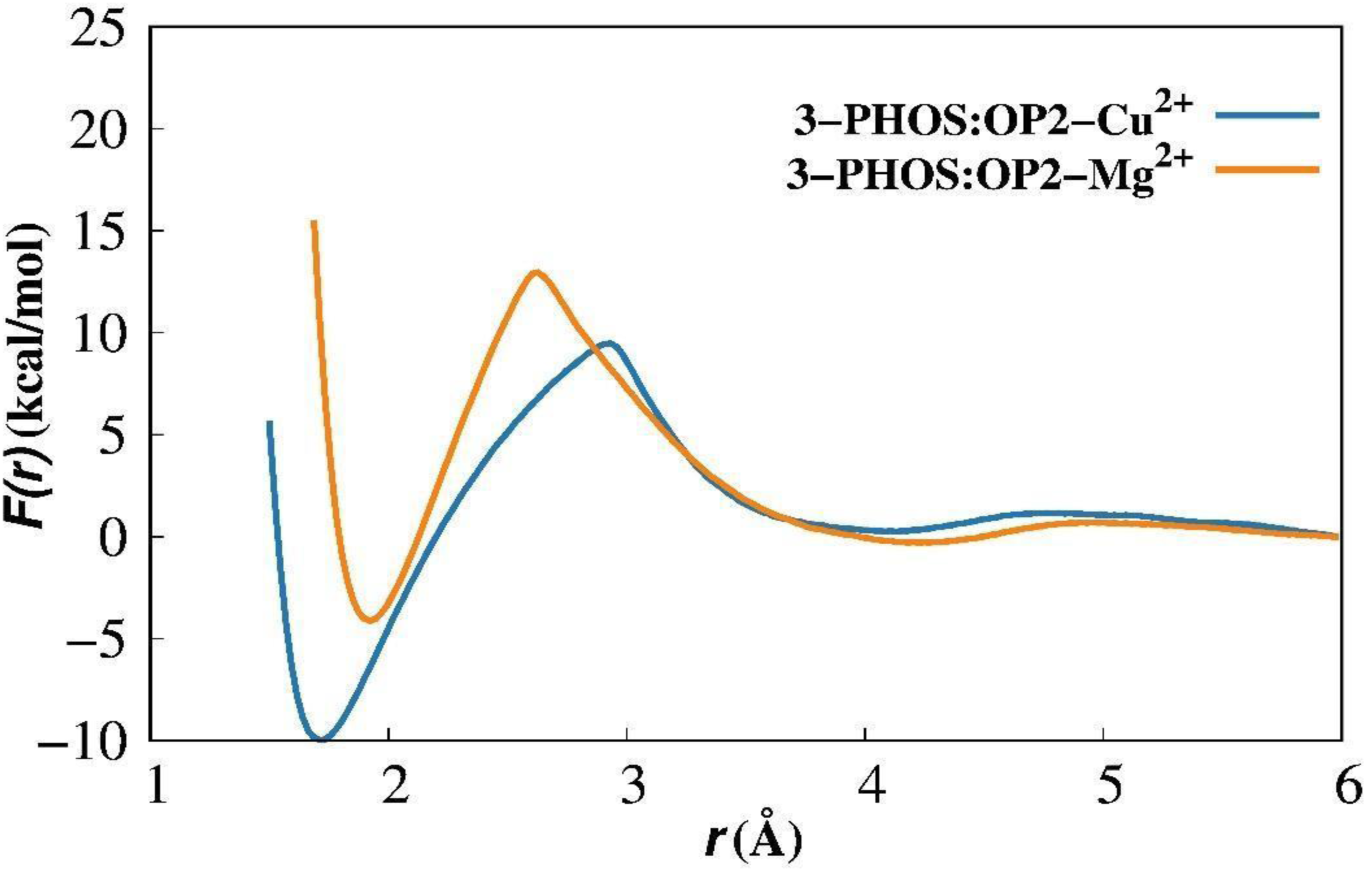
The potential of mean force, F(r), of Cu^2+^ and Mg^2+^ ion with the OP2 atom of the phosphate group of the third nucleotide (cytosine) in the DNA sequence.

Figure 7 shows the *F*(*r*) of Cu^2+^ and Mg^2+^ ions with N7 and O6 atoms of the guanine base in the major grooves and with N3 and O2 atoms of thymine base of the minor grooves. For *F*(*r*) calculations, guanine is specifically chosen because Cu^2+^ ions have been experimentally found to have the highest affinity for the N7 and O6 atoms of guanine base^28^. The results show that *F*(*r*) nearly zero at long separation (6 Å). But as ion comes closes to the binding site, *F*(*r*) decreases and reaches to a minimum ≈ at 4 Å. On further reducing the separation, *F*(*r*) increases and shows a positive hump or maximum around 3 Å. Thereafter, *F*(*r*) decreases slightly with decreasing separation and then shows a sharp increase below 2 Å due to van der Waals repulsion. In contrast to the interaction at the phosphate group, the value of *F*(*r*) in the 1.5 – 3.5 Å region is positive, which suggests that the direct binding of both Cu^2+^ and Mg^2+^ ions in the groove region is energetically unfavorable. Ions are interacting with the electronegative sites of bases in the indirect mode, i.e., with the complete first hydration shell. Unlike experiments, direct binding (complexation or chelation) of Cu^2+^ ions with the electronegative atoms of DNA bases are not observed because of minor structural changes in the DNA. Under the given structural scenario of DNA, the positive *F*(*r*) in the 1.5 – 3.5 Å region could be due to the high energy cost of removing water from the groove region.

**Figure 7.**
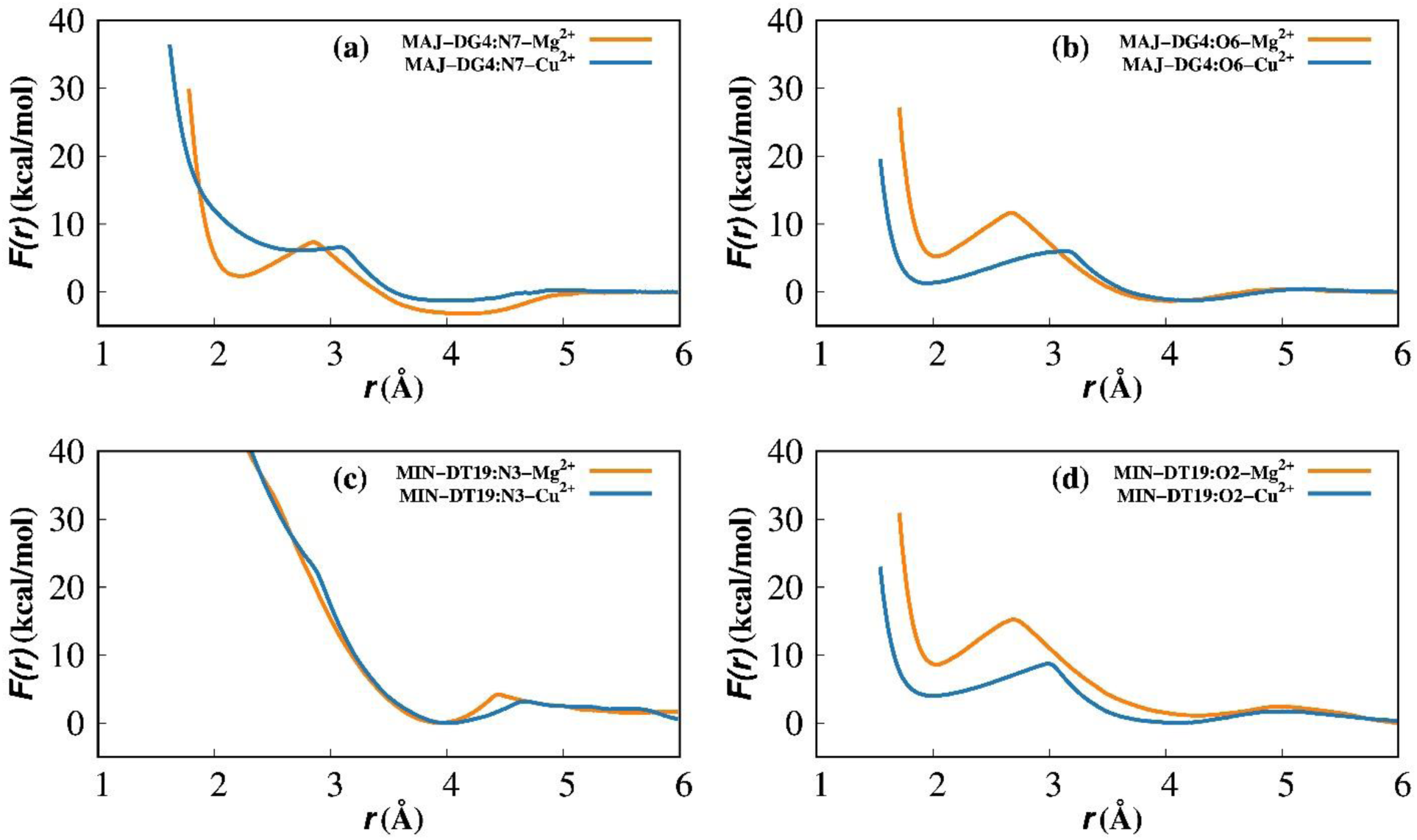
Potential of mean force ( *F*(*r*)) of Cu²⁺ and Mg²⁺ ions with the N7 and O6 atoms of the fourth guanine residue in the major groove, and with the N3 and O2 atoms of the nineteenth thymine residue in the minor groove.

Figure 8 shows the change in the coordination number, *N_c_* of water molecules around the ion as it approaches a given binding site in the DNA in *F*(*r*) calculation, whether at the phosphate group or in the major or minor grooves. It can be clearly seen that above 3 Å *N_c_* is constant, i.e. 6. This implies that above 3 Å ions interact with the DNA in the indirect mode, i.e., with a complete hydration layer. At 3 Å, the *N_c_* of Mg^2+^ remains 6 but becomes 5 for Cu^2+^ ions. Below 3 Å, the coordination numbers of both Cu^2+^ and Mg^2+^ ions are 5, i.e. both ions are interacting with DNA with a penta-coordinated hydration shell.

**Figure 8.**
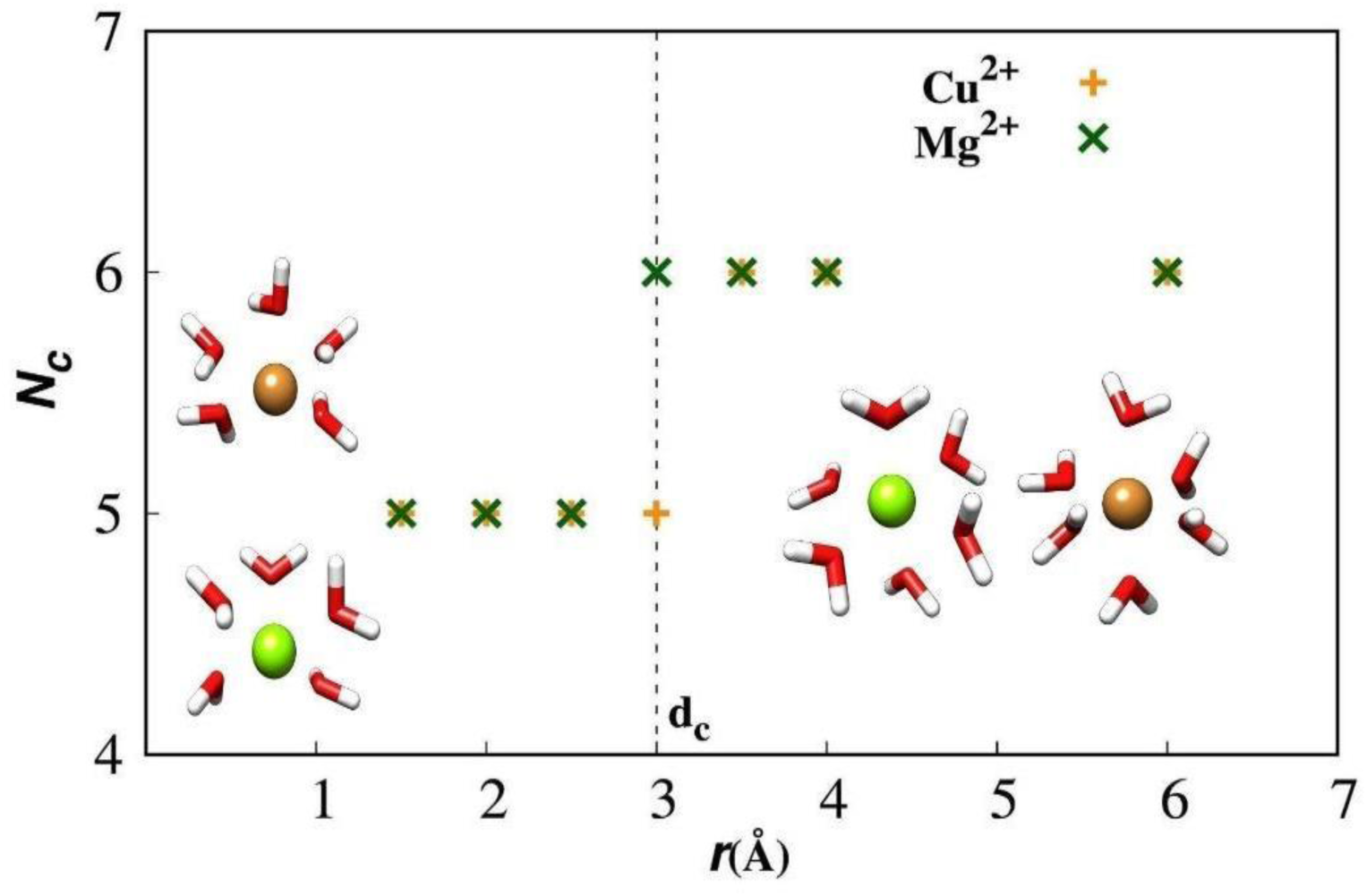
The variation of coordination number, *N_c_*, of water molecules around Cu^2+^ and Mg^2+^ ions as function of distance, i.e., along the reaction coordinate *r* in the *F*(*r*) calculation.

## CONCLUSION

Molecular dynamics simulations were used to study the interactions of Cu^2+^ and Mg^2+^ ions with double-strand DNA. The binding of both Cu^2+^ and Mg^2+^ ions in different parts of DNA, i.e. at phosphate group, in the major and minor grooves were examined through the structure and potential of mean forces (PMFs). Both ions show highest binding affinity in the phosphate group, then in the major groove, and lowest in the minor groove. However, deeper comparisons show that both ions have some notable differences too in the binding behavior. Mg^2+^ ions interact with DNA in the indirect interaction mode, i.e., with complete first hydration shell, consistent with previous simulation and experimental studies. On the contrary, Cu^2+^ ions show labile (variable) coordination shell and thus interact with the DNA through both direct and indirect modes. Owing to the minor structural structure change in the DNA, direct binding of Cu^2+^ ions is observed only at phosphate groups. In direct binding, Cu^2+^ ions interact with DNA through a partial hydration shell (having coordination numbers less than 6) and form direct electrostatic bonds with the oxygens of the phosphate groups. In the ion atmosphere around the DNA, the density of Cu^2+^ ions were found to be higher than that of Mg^2+^ ions. The binding affinity of Cu^2+^ ions at the phosphate group was found to be higher than that of Mg^2+^ ions but lower in the major groove; however, both the ions show similar binding behavior in the minor groove. The distinct binding behavior of similar sized Cu^2+^ and Mg^2+^ ions stem from notable differences in their hydration layers. As far as DNA structure is concerned, only Cu^2+^ brings about some minor structural changes (fraying effect) in the DNA. The direct binding of Cu^2+^ ions in the major and minor grooves with DNA bases are not observed, which is thought to be due to minor structural change in the DNA. It would be interesting to study the binding of Cu^2+^ ions to DNA by applying a bias force on the DNA structure to get an understanding of their direct binding with the bases in the grooves. Overall, the study offers significant insights into the binding behavior of Cu^2+^ ions to DNA, which can be useful for experiments.

## Supporting information

https://drive.google.com/file/d/1aIg9KcPVrvfZe6hTOnlRIb6XXQ4EMEas/view?usp=drive_link

## ACKNOWLEDGMENTS

AS thanks Jawaharlal Nehru University (JNU) for financial support. HOSY thanks Science and Engineering Research Board (SERB) for financial support under project RJF/2021000064. This work was also partially supported by grants from DBT (BT/PR/40251/BITS/137/11/2021) awarded to the Centre for Computational Biology and Bioinformatics, Jawaharlal Nehru University and the MATRICS grant from SERB (MTR/2021/000365) awarded to P.B. The authors are thankful to Dr. Dhananjay Bhattacharyya for insightful discussions.

## DISCLOSURE STATEMENT

The authors report there are no competing interests to declare.

## Notes

### Competing Interest Statement

The authors have declared no competing interest.

