## Supplementary material for "Understanding Cu^+2^ binding with DNA: A molecular dynamics study comparing Cu^2+^ and Mg^2+^ binding to the Dickerson DNA": https://drive.google.com/file/d/1aIg9KcPVrvfZe6hTOnlRIb6XXQ4EMEas/view?usp=drive_link

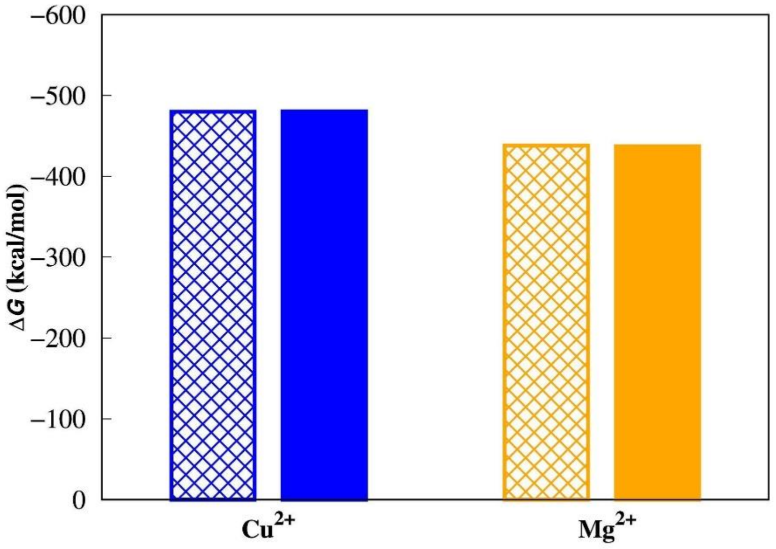


Figure S1. The hydration free energy, $\Delta G$, of $Cu^{2+}$ and $Mg^{2+}$ ions at 300 K. The patterned and solid bars show the experimental and simulation data, respectively. The data are taken from Ref [S1, S2]

### Structural Change of DNA

Ion binding can cause a structural change in the DNA. The structural changes of DNA were characterized by determining the root mean square deviation (RMSD) and the width of the major and minor grooves of DNA. In the RMSD calculation, all heavy atoms of DNA were taken into account and the trajectory of DNA was fitted to the experimental (native) structure. It measures how much the DNA structure changes with respect to the experimental structure upon interaction with the ions. The groove width is computed as the distance between the two opposite phosphorous (P) atoms of both DNA strands in the respective grooves[S3]. For a given groove, distances are averaged over all phosphorus-pairs. Figure 4 shows the variation of the RMSD and the width of the major and minor grooves as a function of time of DNA interacting with both $Mg^{2+}$ and $Cu^{2+}$ ions. The result shows that RMSD of DNA is low. The average value is less than 2.5 Å for both ions. In the interaction of DNA with $Mg^{2+}$ ions, RMSD data simply fluctuates about a mean value suggesting no conformational change in the DNA. However, in the interaction of DNA with $Cu^{2+}$

ions, a minor fraying is seen in the DNA structure between 400 – 700 ns. This suggests that only $Cu^{2+}$ ions cause some minor structural change in the DNA, not $Mg^{2+}$ ions. The variation of groove width provides the same conclusion.

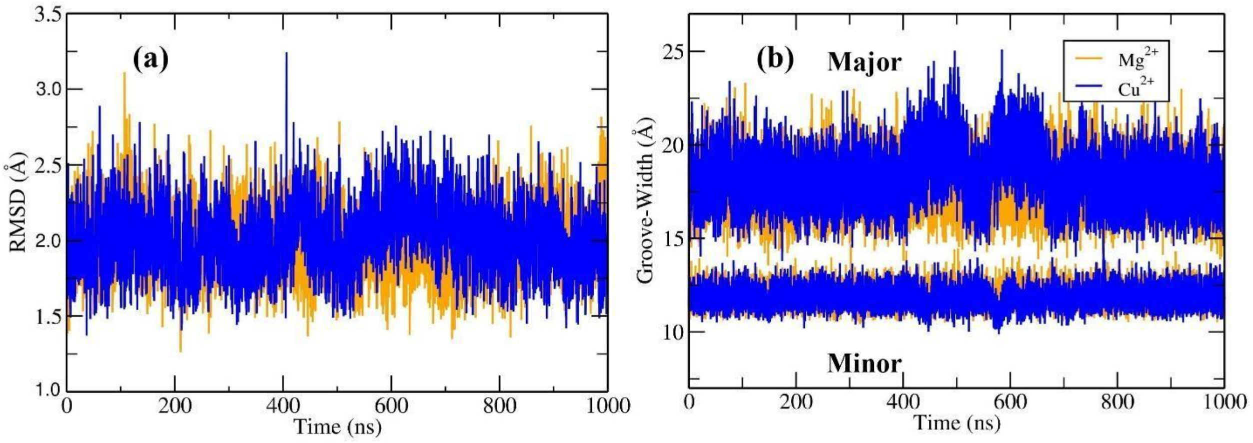


Figure S2. (a) Root mean square deviation and (b) width of major and minor grooves of the DNA.

#### Direct Electrostatic Interaction of $Cu^{2+}$ and $Mg^{2+}$ DNA

Electrostatic interaction can play an important role in ion binding to DNA. Therefore, we analyzed the direct electrostatic interactions of ions with a given electronegative atom of the DNA. Electrostatic interaction was calculated simply using the Coulomb potential. For a given site, the interactions over all $Cu^{2+}$ or $Mg^{2+}$ ions were summed up and then divided by the total number of ions present in the system. Therefore, the $E_c$ data shown in Fig. 3 represents the direct electrostatic interaction of an ion with a given site in DNA. The data is shown for all OP1, N7, and O6 sites. One can see that the data is almost symmetric from the middle. This is because the first half of the figure (residue numbers 1 – 12) represents the first strand and the second half (residues 13 – 24) represents the second strand of the DNA. For any given site, $E_c$ is in general more negative for the $Cu^{2+}$ ion, which suggests that the interaction of $Cu^{2+}$ with DNA is more favorable than that of the $Mg^{2+}$ ion. Although $E_c$ is more favorable for Cu2+ ion, the overall binding affinity would depend on other factors too such as van der Waals interaction, solvation entropy of the binding site in DNA and ion.

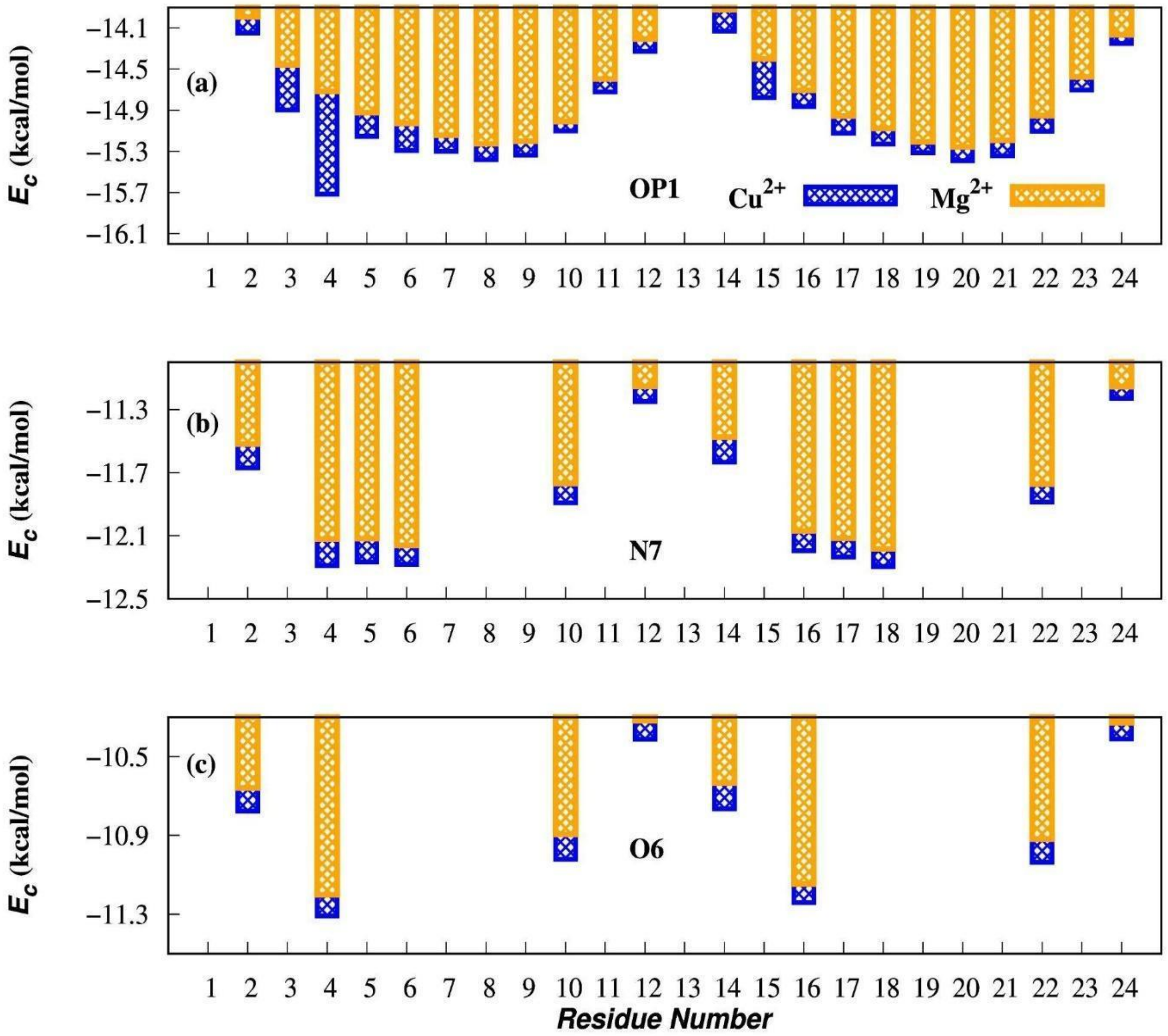


Figure S3. Direct electrostatic interaction, $E_c$, of $Cu^{2+}$ (blue bar) and $Mg^{2+}$ (orange bar) ions with different electronegative sites in DNA. The data is shown as a function of residue number.

### Occupancy of $Cu^{2+}$ and $Mg^{2+}$ around DNA

Ion binding in the different parts of the DNA was also investigated in terms of percentage occupancy, which was defined as follows,

$$\%\ occupancy = \frac{Number\ of\ ions\ within\ 5\ \text{Å}\ of\ all\ electronegative\ atoms\ in\ the\ groove}{Total\ number\ of\ ions\ within\ 5\ \text{Å}\ of\ DNA}$$

where 5 Å cutoff distance was chosen based on $g(r)$. This is the distance that includes the first peak of the $g(r)$ corresponding to the ions indirectly interacting with the DNA. The percentage occupancy provides a measure of the binding affinity of ions for DNA. Fig. 4(a) displays the percentage of ions present within 5 Å of DNA, i.e. it is the ratio of the number of ions present

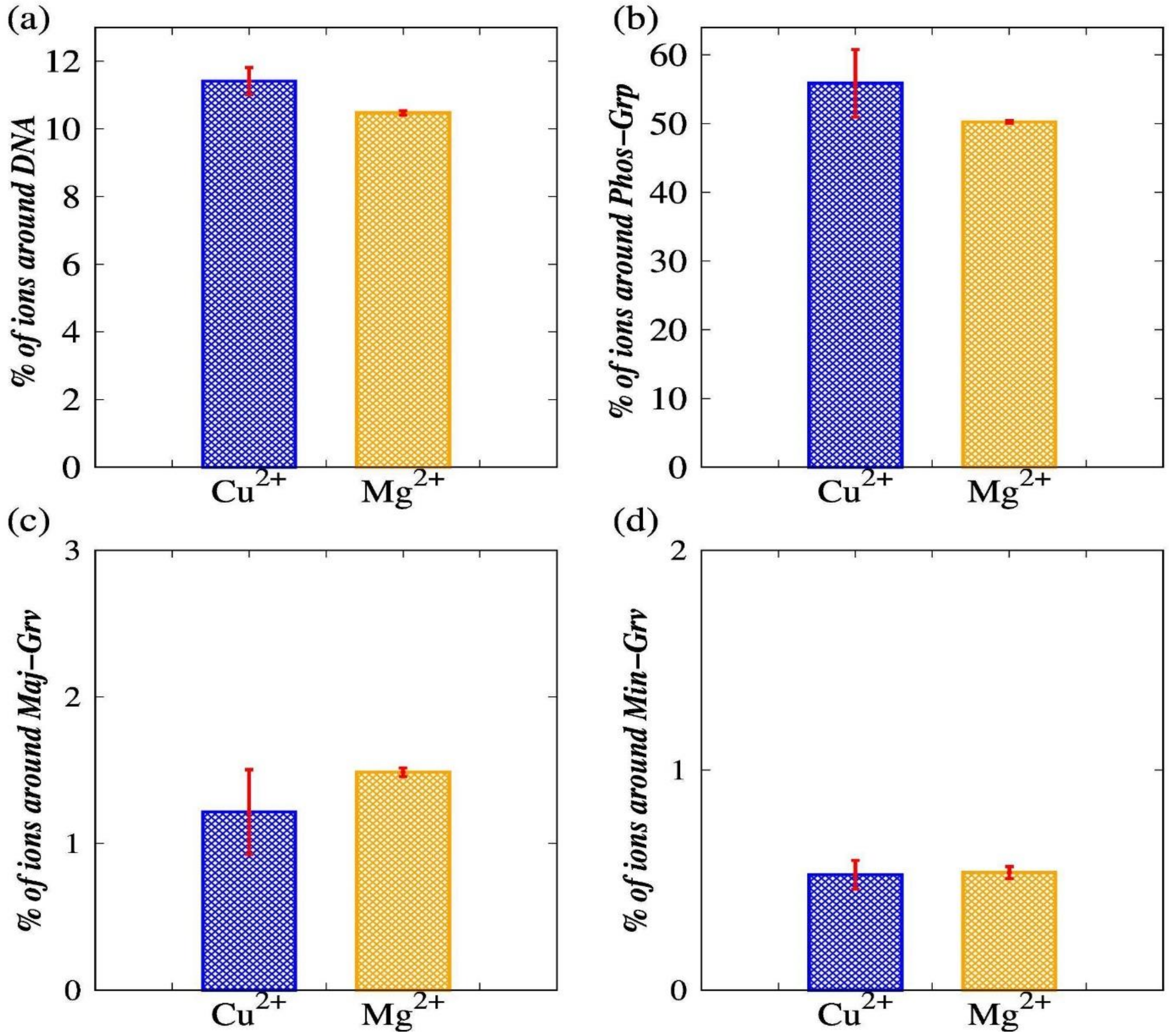


Figure S4. (a) The percentage of ions within 5 Å of whole DNA. The percentage occupancy at the (b) phosphate group, (c) major groove, and (d) minor groove.

within 5 Å of the DNA to the total number of ions present in the system (82 ions). The percentage occupancy shown in Fig. 4(b) – 4(d) are calculated using the above equation. In other words, Fig. 4(b) – 4(d) are partitions of Fig. 4(a). The result shown in Fig. 4(a) clearly shows that the number of $Cu^{2+}$ ions within 5 Å of the DNA is higher than that of $Mg^{2+}$ ions. This suggests that $Cu^{2+}$ ions have greater affinity for DNA than $Mg^{2+}$ ions. Figures 4(b) – 4(d) show that for both ions, the highest binding affinity is in the phosphate group, followed by the major groove, then lowest in the minor groove. The comparison indicates that $Cu^{2+}$ ions have a higher binding affinity for the phosphate groups, while $Mg^{2+}$ ions have a higher affinity for the major groove of DNA. In contrast, the binding behavior of $Cu^{2+}$ and $Mg^{2+}$ ions in the minor groove are similar. The $g(r)$ analyses provide the same predictions.

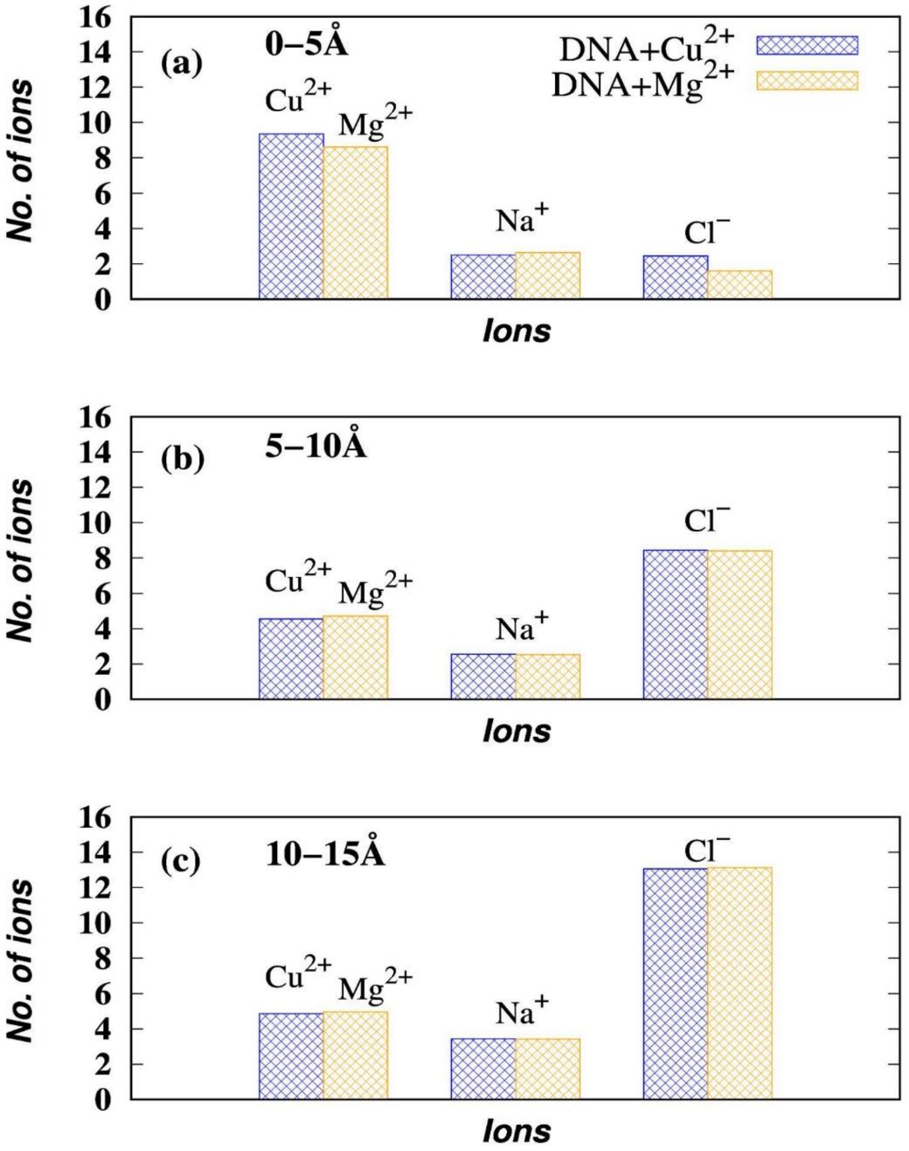


Figure S5. The counts of divalent ($Cu^{2+}$ and $Mg^{2}$) and background ($Na^{+}$, and $Cl^{-}$) ions in different distance intervals (a) 0 – 5 Å, (b) 5 – 10 Å, and (c) 10 – 15 Å around DNA.

#### Counts of Ions in Different Distance Intervals around DNA

The ion distribution around DNA changes with distance. As shown in Figure 5(a), within a 5 Å radius of the DNA, divalent ions are more abundant than $Na^+$. This happens because DNA carries 22 units of negative charge and the electrostatic interaction between the negative charges of the DNA and positive divalent ions is more as compared to the Na+ ions. Since DNA is negatively charged, due to repulsion the $Cl^-$ ions around DNA within 5Å would be very less. In Fig. 5(c) and (d), the number of $Cl^-$ ions is increasing and the number of divalent ions is decreasing. This happens because the negative charge of the DNA gets screened by the positive ions which are present in the close proximity of the DNA. The ionic environment around DNA is now positively charged, it will start attracting negative ions present in the system. We found that in close proximity to DNA, there were more $Cl^-$ ions present when $Cu^{2+}$ ions interacted with DNA compared to when $Mg^{2+}$ ions interacted with DNA.
